# The Limited Range of Motion of the Knee Does Not Fully Explain the Altered Neural Control of Plantar Flexors During Gait in Non-Neurological Knee Flexion Contracture

**DOI:** 10.64898/2026.06.08.730170

**Authors:** Carlos Cruz-Montecinos, Tjeerd W. Boonstra, Huub Maas

## Abstract

Knee flexion contracture (KFC) may occur in the late stages of arthropathies, including osteoarthritis and haemophilic arthropathy. The impact of KFC on neuromuscular control remains unclear, particularly for the affected ankle plantar flexors. Surface electromyography (EMG) is widely used to assess muscle activation patterns, whereas intermuscular (EMG–EMG) coherence provides insight into common neural input. In this study we compared the neural control of ankle plantar flexors during gait between individuals with haemophilia and KFC (chronic; n = 8), and healthy individuals without (control; n = 15) and with an artificial constraint (artificial; n = 15). Bipolar EMG from plantar flexors was recorded during 30-m overground walking (1 m/s). Intermuscular coherence was estimated at 8–60 Hz during the stance phase and significance was determined using a permutation method. The chronic group showed greater knee flexion than controls (24–29 deg vs 2–20 deg), higher EMG amplitude at foot contact, and increased intermuscular coherence in the alpha (8–12 Hz) and beta (12–30 Hz) bands at mid-stance. Despite comparable sagittal knee kinematics between constrained conditions (chronic: 24–29 deg; artificial: 20–32 deg), early-stance EMG amplitude and mid-stance beta-band intermuscular coherence were higher in the chronic group across plantar-flexor pairs. Increased plantar-flexor activation in the chronic group suggests higher neural drive, while higher intermuscular coherence reflects greater common input to the plantar flexors. These findings indicate that limited ROM alone does not explain the altered neural control of plantar flexors, suggesting neural adaptations associated with non-neurological chronic KFC.

## Introduction

Knee flexion contracture (KFC) is common in neurological conditions (e.g., cerebral palsy), but may also occur in the late stages of arthropathies, including osteoarthritis, rheumatoid arthritis, and haemophilic arthropathy (Lee et al., 2022; Marron et al., 2025). A knee flexion contracture implies a mechanical block in obtaining full knee extension (Marfo et al., 2018). In people with KFC, the joint constraint is accompanied by pain, worse functionality and increased motor demand (Campbell and McGonagle, 2021; Murphy et al., 2014). Flexion contractures affect gait because the limb cannot fully extend to support the body, leading to compensatory changes in ankle and foot mechanics particularly during stance (Michael Attias et al., 2016). Non-neurological KFC leads to long-term functional limitations (Campbell and McGonagle, 2021), affecting about one-third of patients with knee osteoarthritis (Campbell and McGonagle, 2021; Ritter et al., 2007).

In chronic KFC, the primary treatment approach to restore the range of motion (ROM) is orthopedic surgery, such as total knee replacement, combined with surgical correction and release of contracted soft tissues (Kim et al., 2017; Scuderi and Kochhar, 2007). Although total knee replacement has been reported to be an effective treatment for people with osteoarthritis (Kahlenberg et al., 2018), reducing the KFC remains a particular challenge, especially in cases of a long-term history of contracture or severe deformity (Sappey-Marinier et al., 2024). Current surgical approaches mainly address the mechanical components of KFC (Sappey-Marinier et al., 2024; Zhai et al., 2019). This does not account for potential long-term central nervous system adaptations that may influence gait recovery (Cruz-Montecinos et al., 2021). Because KFC alters limb posture during stance and increases the demand for maintaining body support and forward progression, muscles acting across adjacent joints may also adapt their control strategies (Michael Attias et al., 2016; Cerny et al., 1994). In this context, the ankle plantar flexors are especially relevant because they contribute to body support and forward progression during the stance phase of gait (Meinders et al., 1998; Neptune et al., 2001). Determining whether these locomotion adaptations arise from joint mechanics or from persistent motor control changes is clinically important, as it may help guide rehabilitation and orthopedic interventions from a sole focus on mechanical factors toward incorporating neuromechanical considerations.

Surface electromyography (EMG) is commonly used to examine the magnitude and timing of muscle activation during gait (Mills et al., 2013). Since the amplitude and frequency of the EMG signal are shaped by motor unit firing behavior, they serve as an indirect index of neural drive (Farina et al., 2010). Based on this, intermuscular (EMG–EMG) coherence provides a complementary measure of the shared presynaptic drive to synergistic motor neuron pools (Boonstra et al., 2009b; Farmer et al., 1993). The frequencies above 20 Hz (i.e., beta and gamma bands) are traditionally thought to reflect descending corticospinal drive (Brown et al., 1999; Hansen et al., 2005). Recent evidence also underscores their increased power during proactive (e.g., obstacle negotiation) and proprioceptive (e.g., perturbation-based) locomotor tasks to maintain stability (De Freitas et al., 2026). Coherence in lower frequencies (alpha, 8–12 Hz) may capture subcortical or reticulospinal contributions, potentially reflecting compensatory neural strategies or automated postural adjustments during complex gait (De Freitas et al., 2026).

In individuals with non-neurological KFC, a critical unresolved question remains whether observed neural alterations reflect a primary reorganization of central drive or are secondary to mechanical joint restrictions (Cruz-Montecinos et al., 2021; Spomer et al., 2022). Distinguishing between mechanical and neural adaptations requires a design that separates acute mechanical restriction from chronic impairment. For that, simulation of KFC has been used as a model to understand the acute effects on kinematics and muscle activity (M. Attias et al., 2016; Cerny et al., 1994; Harato et al., 2008).

Accordingly, we addressed two questions: *(1) How is neural drive to the ankle plantar flexors altered in individuals with chronic knee flexion contracture? and (2) To what extent is this altered neural drive explained by the mechanical constraint at the knee?* To answer these questions, we compared the neural control of the ankle plantar flexors during gait in individuals with haemophilia, and KFC with that in healthy individuals walking without and with an imposed knee constraint.

## Methods

### Study design

This study employed a cross-sectional comparative design involving three gait conditions. The first condition included individuals with haemophilia and chronic KFC (chronic). People with haemophilia have a genetic bleeding disorder characterized by a deficiency of coagulation factors VIII or IX (Berntorp et al., 2021). They were selected because they may experience advanced arthropathy due to repeated bleeding episodes, which lead to synovial inflammation and irreversible cartilage damage (Berntorp et al., 2021). These changes may be accompanied by excessive fibrous or scar tissue within a joint, leading to restricted ROM, and often result in severe joint contracture at middle age (Silva and Luck, 2007).

The second group included healthy adults walking without a knee joint constraint (control). The third included healthy adults walking with an imposed knee restriction intended to simulate a mechanically constrained knee (artificial). Some of the data used have been reported in a previous study that addressed a different research question related to comparing acute versus chronic knee constraints on muscle synergies (Cruz-Montecinos et al., 2021). The primary outcomes were stance-phase kinematics, plantar-flexor EMG amplitude, and intermuscular coherence between plantar-flexor muscles. This study was approved by the local ethical committee at Metropolitan Health Service and conducted in agreement with the Declaration of Helsinki. All participants were informed about the purpose and procedures of the project and gave their written informed consent to participate in the study.

### Participants

Two groups were analyzed: a chronic (n = 8) and a healthy group (control; n = 15) tested without constraint and with an artificial KFC (artificial; n=15). For characteristics of participants see Table 1. Participants in the chronic KFC group were selected based on a clinical diagnosis of knee flexion contracture diagnosed by a medical doctor specialized in hemophilic arthropathy, less than 30 degrees of knee ROM, over 18-years of age and under 65-years, prophylaxis treatment with deficient factor (i.e., VIII or IX), and body mass index less than 30. Exclusion criteria included history of hip, knee or ankle arthroplasty, equinus foot, inability to walk without an assistive device (e.g., walker, cane), history of muscle or joint bleeding in lower limbs in the last two-months, chronic cardiac and/or respiratory pathology and neurological disease.

**Table 1.** Characteristics of control group and people with non-neurological knee flexion contracture (Chronic).

| Clinical Characteristic | Control (n=15) | Chronic (n=8) | p-value |
| --- | --- | --- | --- |
| Age (years) | 31.5 ± 10 | 39.1 ± 16 | 0.167 |
| Body mass index | 24.7 ± 1.9 | 26.4 ± 2 | 0.342 |
| Pain during walking (VAS 0-10) | 0 [0 0] | 4.0 [1 8] | **<0.001** |
| Chronicity (years) | NA | 15 [10 20] | **NA** |

Parametric distribution: Mean ± SD. Nonparametric distribution: Median [Range]. VAS, Visual Analogue Scale. NA, not applicable. Significant differences (p<0.05) are in bold.

The control group consisted of healthy males (18–65 years), body mass index less than 30, with no history of haemophilia or neuromuscular and musculoskeletal disorders affecting locomotor patterns.

### Experimental procedure

Walking trials were standardized across groups. Each participant completed the protocol by walking barefoot over a 30-meter walkway corridor at a fixed speed of 1 m/s, twice, under standardized conditions. Before EMG recording, each participant practiced twice at the selected speed. The use of a fixed walking speed reduced between-subject variability associated with self-selected pace and enabled a more consistent comparison of stance-phase muscle behavior across conditions. In the artificial constraint condition, healthy participants walked with an externally imposed knee restriction that acutely limited knee ROM to 20 deg–40 deg **(Figure 1)**. This level of restriction was selected to approximate the passive ROM limitation observed in the chronic KFC group. Each participant practiced with the external device three times for 10 meters. Ten gait cycles from the middle of each walking trial were selected for the final analysis, yielding 20 cycles in total per participant. Initial contact was detected using a pressure sensor positioned under the heel. The analysis focused on the stance phase (first 60% of gait cycle), since it includes load acceptance, body support, and forward movement—functions where the plantar flexors play a key mechanical and neural role (Neptune et al., 2001). Time-resolved analysis, time-normalized to 100% of the stance phase, allowed for comparisons of plantar-flexor activation profiles, knee kinematics, and intermuscular coherence across conditions.

**Figure 1.**
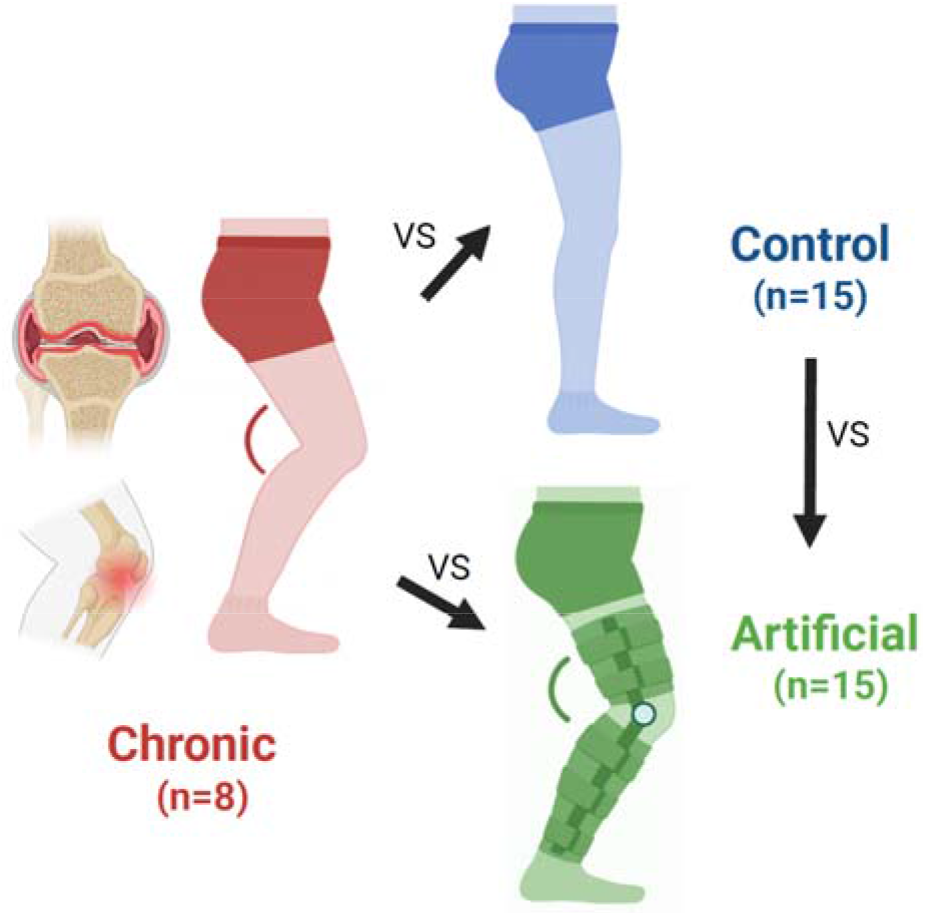
Experimental design and study groups. Individuals with chronic knee flexion contracture (Chronic; red; n = 8) associated with haemophilic arthropathy were compared with healthy participants walking under two conditions: unconstrained walking (Control; blue; n = 15) and walking with an externally imposed knee flexion restriction (Artificial; green; n = 15).

### EMG acquisition

Bipolar surface EMG was recorded from the medial gastrocnemius (MG), lateral gastrocnemius (LG), and soleus (SOL) of the affected or tested limb during walking. Surface EMG activity was recorded using pa rs of Ag-AgCl electrodes (Kendall H124SG) placed over each muscle with an inter-electrode distance of 2 cm. The most affected limb was analyzed in the chronic group, whereas the dominant limb was analyzed in the healthy participants. Signals were collected using a wireless EMG system (MyoSystem DTS, Noraxon USA Inc., Scottsdale, AZ, USA) at a sampling frequency of 1500 Hz.

The MG, LG, and SOL muscles were selected because they act as synergistic ankle plantar flexors and are strongly involved in stance-phase support and propulsion. Skin preparation and electrode placement were based on anatomical reference points as indicated in established guidelines and current recommendations (Besomi et al., 2020; Hermens et al., 2000).

### Kinematic assessment

Sagittal-plane knee flexion-extension kinematics were obtained using inertial measurement units (IMUs; Xsens, Enschede, the Netherlands) placed on the thigh and shank. The sensors were synchronized with the EMG and pressure-sensor signals and sampled at 75 Hz. Sagittal joint position was referenced to upright standing using magnetometer information. Knee flexion-extension was then estimated from the relative orientation of the thigh during gait (Cruz-Montecinos et al., 2021). The resulting kinematic signals were low-pass filtered at 10 Hz and segmented between consecutive heel strikes. For the kinematic analysis, data from one participant in the control group was excluded because of signal loss.

### EMG data analysis

Surface EMG signals were processed offline using a custom MATLAB routine. Before coherence analysis, EMG signals were band-pass filtered between 20 and 500 Hz using a sixth-order Butterworth filter applied in forward and reverse directions and full-wave rectified. The rectified EMG were then submitted to a phase-based demodulation procedure in which each signal was mean-centered, transformed into its analytic representation using the Hilbert transform, and reduced to a phase-only signal by taking the cosine of the instantaneous phase (Boonstra et al., 2009a). This procedure was used to reduce artefacts in coherence estimates associated with rapid changes in EMG amplitude during gait. Spectral estimates were obtained across the stance phase, time-normalized from 0 to 100%, at 101 equally spaced time points using a 500-ms analysis window.

Intermuscular coherence was estimated between plantar-flexor muscle pairs between MG, LG, and SOL using a multivariate event-related spectral decomposition approach implemented in MATLAB. The function used to compute time-frequency coherence between pairs of continuous signals is available at (https://github.com/tjeerdboonstra/coherence-analysis). The analysis was based on the Fourier decomposition using a sliding window approach with relative (time normalized) rather than an absolute time step. For each point and stride, auto-spectra and cross-spectra were computed after detrending and Hanning windowing, and coherence was derived from the average complex coherency across cycles (Mehrkanoon et al., 2013). Spectral estimates were calculated using zero-padded fast Fourier transforms, and coherence was examined within the 8-60 Hz range. Coherence values within the frequency range of interest were used to derive time-resolved band profiles across the normalized stance phase. In addition to total coherence, a significance-masked coherence measure was obtained by retaining only values exceeding the confidence interval estimated from the number of gait cycles included at an alpha level of 0.05 (Amjad et al., 1997).

### Statistical analysis

For both the SPM and coherence analyses, paired-samples t-tests were used for within-subject comparisons (Control vs Artificial), whereas independent-samples t-tests were used for between-subject comparisons (Control vs Chronic and Artificial vs Chronic).

Time-dependent kinematic and EMG amplitude waveforms were compared using Statistical Parametric Mapping (SPM) (Pataky, 2010). For that, the open-source MATLAB-based spm1d-package was used (http://www.spm1d.org/index.html). For the coherence analysis, time–frequency coherence maps were statistically assessed using a custom cluster-based permutation framework following the principles described by Maris and Oostenveld (Maris and Oostenveld, 2007). Coherence values were Fisher z-transformed (Laine et al., 2014; Rosenberg et al., 1989), and Welch’s t-statistic maps were generated for each comparison. Adjacent supra-threshold time–frequency bins were grouped into clusters, and significance was determined against the permutation distribution of the maximum cluster statistic obtained from 4000 permutations. For within-subject comparisons, condition labels were randomly swapped within each participant, whereas for between-subject comparisons, group labels were randomly permuted across participants. This procedure controlled the family-wise error rate across the time–frequency plane (Maris and Oostenveld, 2007). Statistical significance was set at p < 0.05.

## Results

### Kinematic differences during stance

SPM analysis showed significant differences between the chronic KFC and healthy control groups, as well as between the artificial constraint and healthy control conditions, for the whole stance phase (both p < 0.001). *Chronic vs Control*. The chronic group walked with markedly greater knee flexion angle and less ROM (mean between 24 deg and 29 deg) during the whole stance phase than the healthy control group (mean between 2 deg and 20 deg). *Chronic vs Artificial*. No significant differences were found between the artificial KFC and chronic KFC groups, indicating that the artificial KFC replicates the knee movement pattern seen in the chronic condition group. *Artificial vs Control*. The artificial KFC condition resulted in a knee flexion pattern with greater knee flexion and angle and smaller ROM (mean; between 20 deg and 32 deg), showing a significant difference compared to control gait during the whole stance phase **(Figure 2 and complementary Figure 1)**.

**Figure 2.**
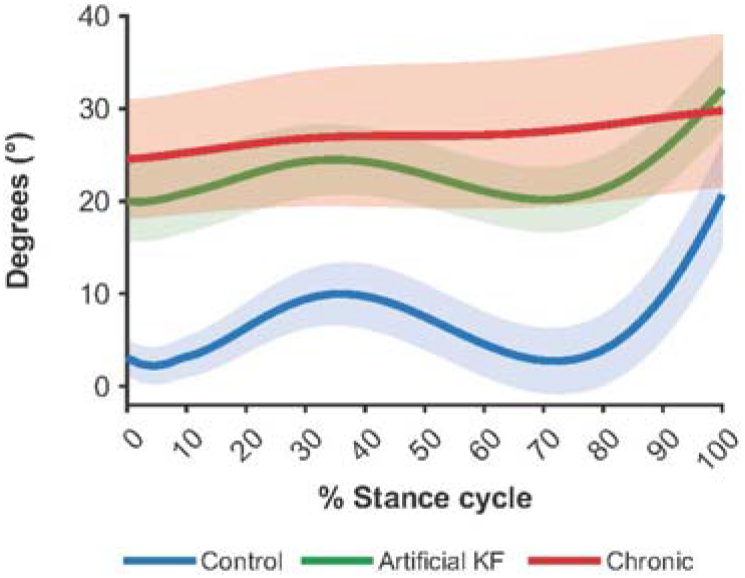
Sagittal-plane knee kinematics during the stance phase of gait in the control (blue; n=14), artificial knee flexion (Artificial; green; n=14), and chronic knee flexion contracture (Chronic; red; n=8) conditions. Data from one participant in the control group was excluded because of signal loss (n=14). Solid lines represent the mean waveform, and shaded areas indicate 95% confidence intervals.

### EMG amplitude patterns during gait

#### Chronic vs Control

Compared with control group, the chronic group showed higher plantar-flexor EMG amplitudes at foot contact and during early stance, followed by lower EMG amplitude during the second half of stance. *Chronic vs Artificial*. Despite broadly comparable joint kinematics, significantly higher early-stance EMG amplitudes were observed in the chronic group compared with the artificial group. *Artificial vs Control*. Compared with the unconstrained control condition, the artificial constraint showed slightly higher plantar-flexor EMG amplitude during early stance, followed by lower EMG amplitude during the late stance-to-swing transition **(Figure 3 and complementary Figure 2)**.

**Figure 3.**
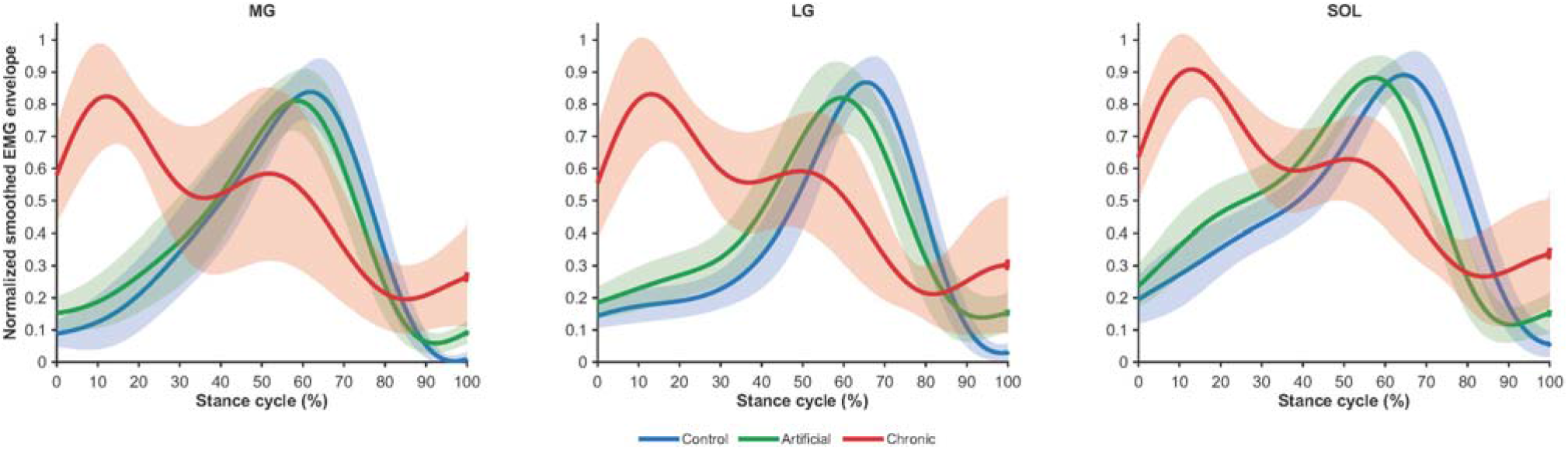
Smoothed plantar-flexor EMG amplitude envelopes during the stance phase for the medial gastrocnemius (MG), lateral gastrocnemius (LG), and soleus (SOL) muscles across the control (n=15; blue;), artificial knee flexion (Artificial; green; n=15), and chronic knee flexion contracture (Chronic; red; n=8) conditions. Solid lines represent the mean waveform, and shaded areas indicate 95% confidence intervals.

### Intermuscular coherence findings

Intermuscular coherence was observed after foot contact (during the double support phase) in the alpha and beta frequency bands, reduced during mid-stance, and started to increase again just before toe-off. This temporal pattern was mainly observed in control and artificial conditions. *Chronic vs Control*. Compared with the healthy control group, the chronic group showed higher coherence in the alpha and beta bands during mid-stance (40-80% of the stance phase) for all plantar-flexor pairs. Additionally, the chronic group showed higher coherence in gamma band between the LG-SOL and MG-LG pairs. *Chronic vs Artificial*. When the chronic group was compared with the artificial condition, higher coherence during mid-stance remained evident in the beta band across all plantar-flexor pairs. Increased alpha-band coherence was observed only for the MG-LG and MG-SOL pairs, whereas increased gamma-band coherence was restricted to the MG-LG pair. *Artificial vs Control*. In the healthy group, artificial constraint condition only showed a higher coherence in beta band between LG-MG just before toe-off (80-100%) **(Figure 4-6)**. Summary results are shown in **Table 2**.

**Table 2.** Summary of intermuscular coherence results.

| Pairs | Bands | Chronic vs Control | Chronic vs Artificial | Artificial vs Control |
| --- | --- | --- | --- | --- |
| MG-LG | Gamma | * | * | --- |
|  | Beta | * | * | * |
|  | Alpha | * | * | --- |
| LG-SOL | Gamma | * | --- | --- |
|  | Beta | * | * | --- |
|  | Alpha | * | --- | --- |
| MG-SOL | Gamma | --- | --- | --- |
|  | Beta | * | * | --- |
|  | Alpha | * | * | --- |

Summary of significant between-group differences in intermuscular coherence across plantar-flexor muscle pairs and frequency bands. (\*) Indicates a significant difference ( $p < 0.05$ ); (---) Indicates no significant difference. Muscle pairs include medial gastrocnemius-lateral gastrocnemius (LG-MG), lateral gastrocnemius-soleus (LG-SOL), and medial gastrocnemius-soleus (MG-SOL).

**Figure 4.**
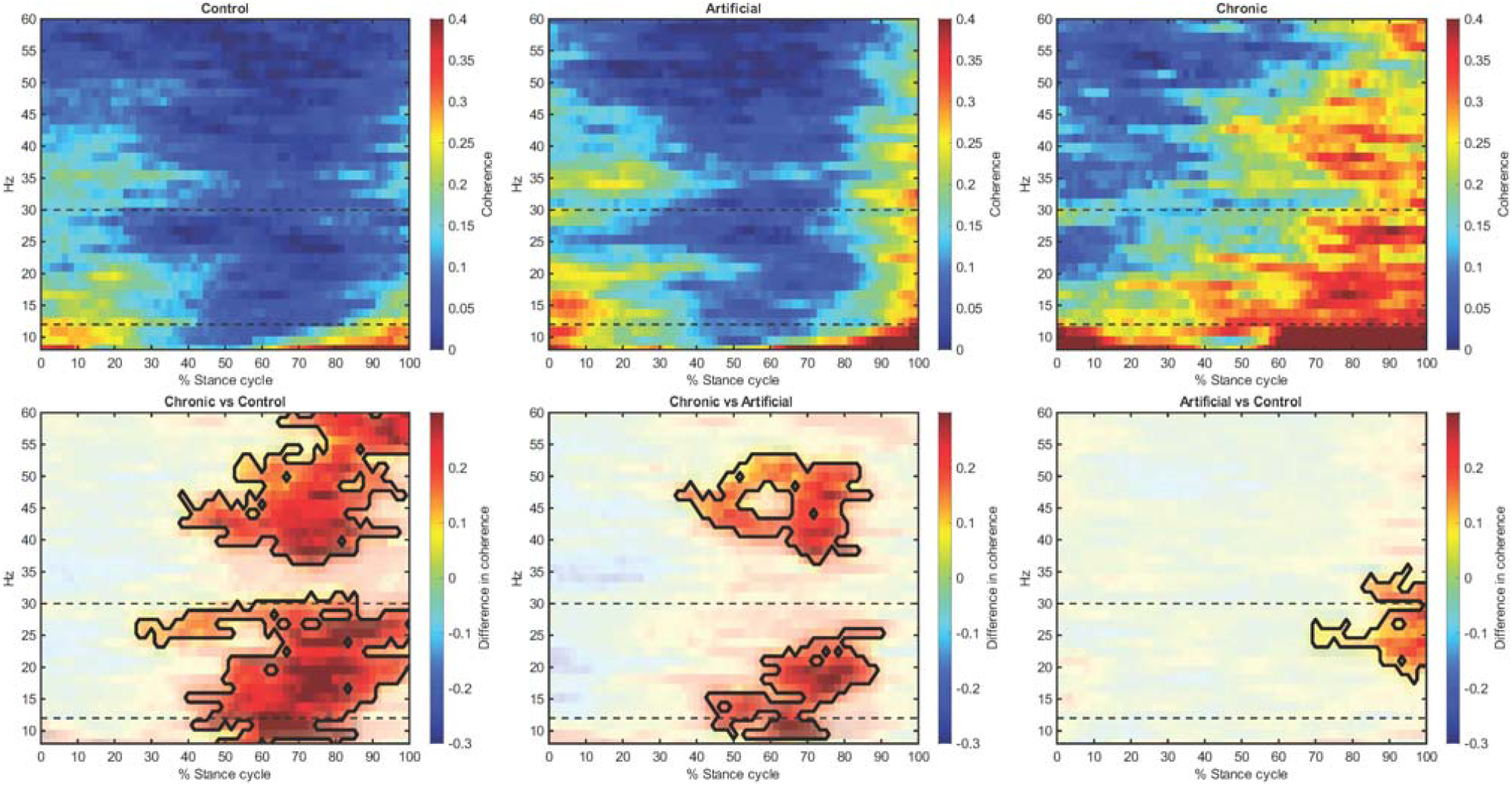
Time-frequency intermuscular coherence between MG-LG during the normalized stance phase (0–100%) in Control (n=15), Artificial (n=15), and Chronic (n=8). Upper panels show coherence maps from 8 to 60 Hz; warmer colors indicate higher coherence. Dashed lines delimit alpha (8–12 Hz), beta (12–30 Hz), and gamma (30–60 Hz) bands. Lower panels show coherence differences between conditions. Black contours indicate significant clusters from cluster-based permutation testing (p < 0.05).

**Figure 5.**
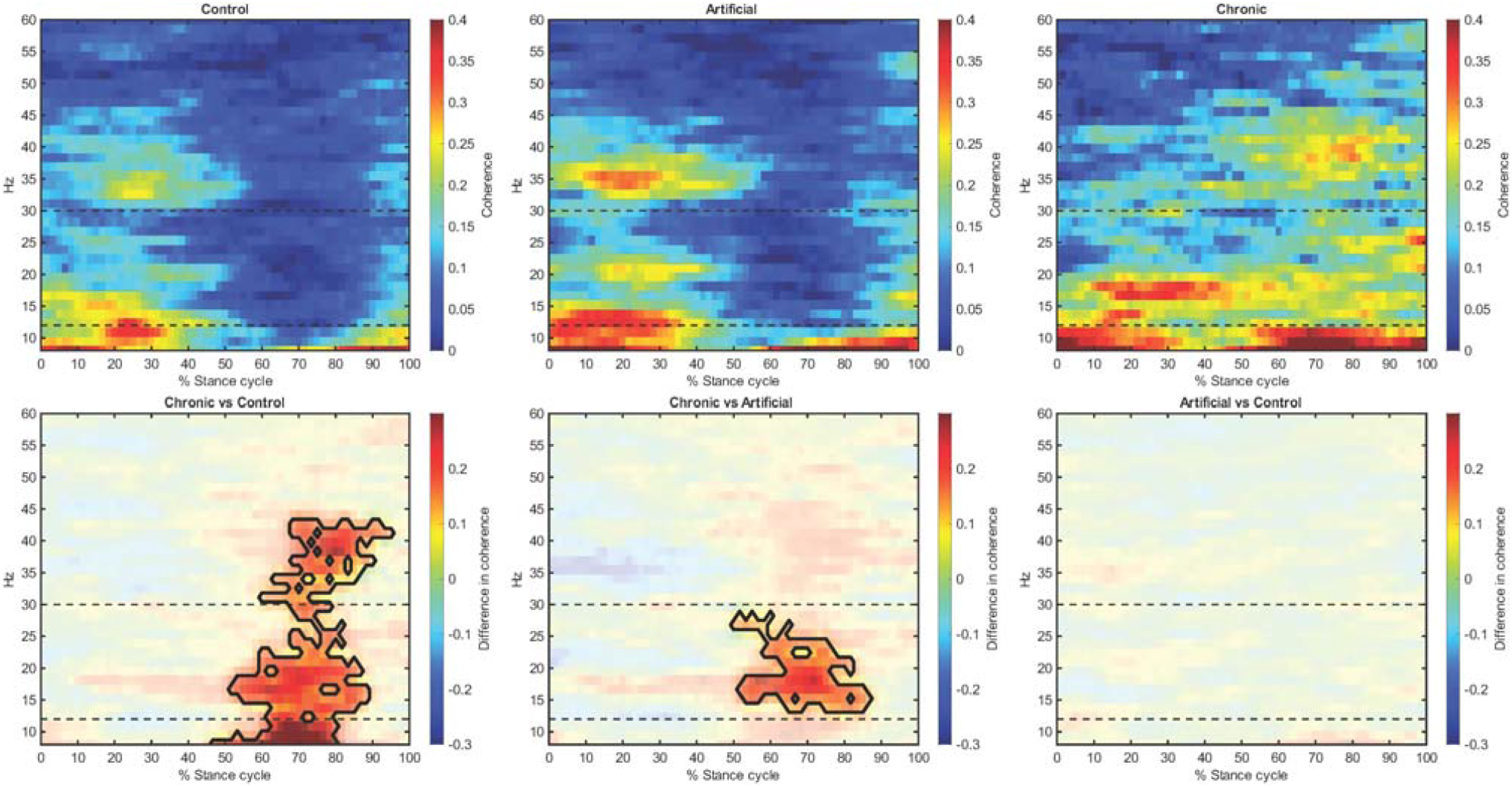
Time-frequency intermuscular coherence between LG-SOL during the normalized stance phase (0–100%) in Control (n=15), Artificial (n=15), and Chronic (n=8). Upper panels show coherence maps from 8 to 60 Hz; warmer colors indicate higher coherence. Dashed lines delimit alpha (8–12 Hz), beta (12–30 Hz), and gamma (30–60 Hz) bands. Lower panels show coherence differences between conditions. Black contours indicate significant clusters from cluster-based permutation testing (p < 0.05).

**Figure 6.**
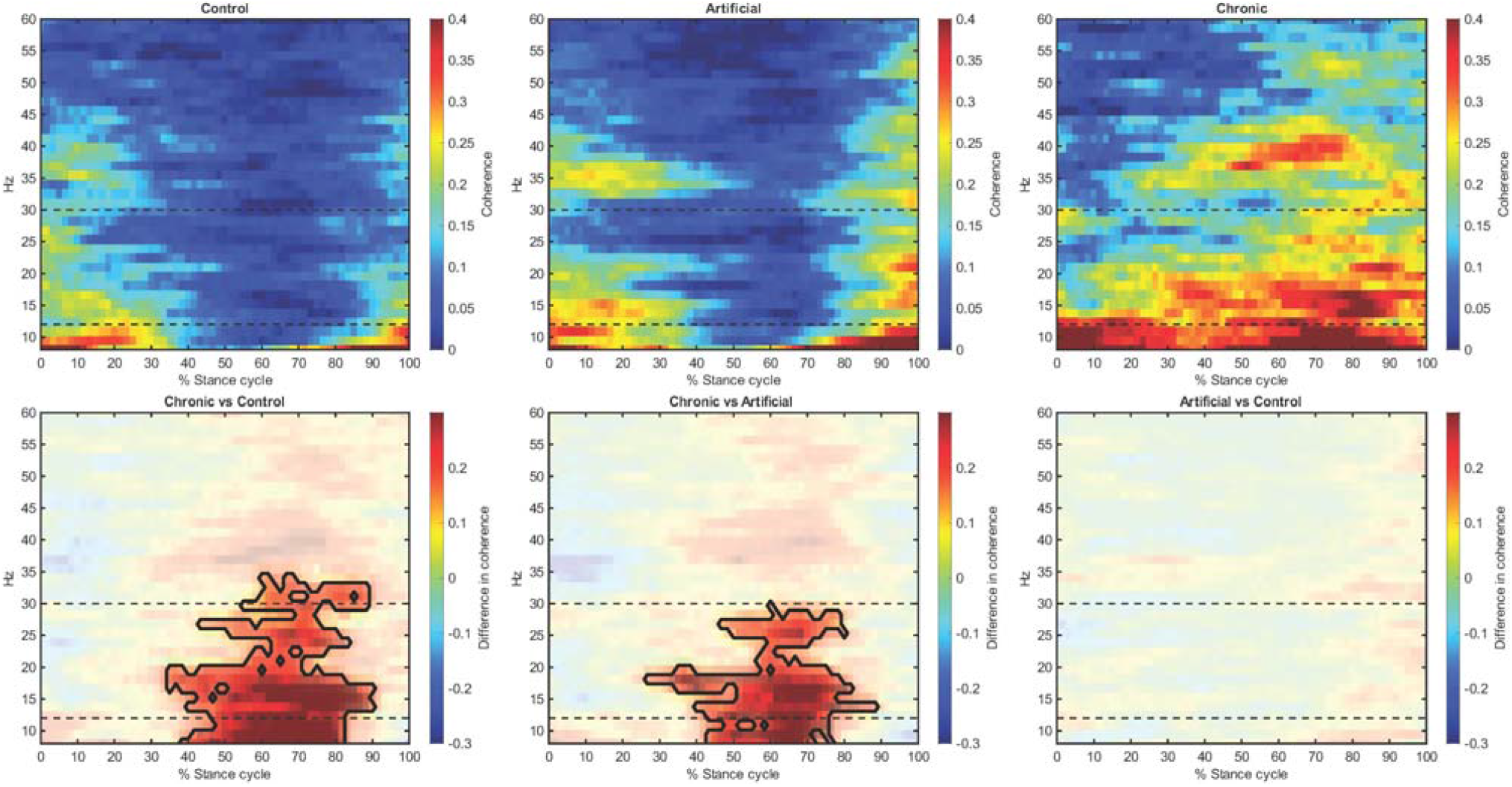
Time-frequency intermuscular coherence between MG-SOL during the normalized stance phase (0–100%) in Control (n=15), Artificial (n=15), and Chronic (n=8). Upper panels show coherence maps from 8 to 60 Hz; warmer colors indicate higher coherence. Dashed lines delimit alpha (8–12 Hz), beta (12–30 Hz), and gamma (30–60 Hz) bands. Lower panels show coherence differences between conditions. Black contours indicate significant clusters from cluster-based permutation testing (p < 0.05).

## Discussion

The aim of our study was to compare the neural control of the ankle plantar flexors during gait in individuals with haemophilia and KFC with that in healthy individuals walking without and with an imposed knee constraint. There were three main findings in this study. First, the chronic group exhibited markedly greater knee flexion angle and less ROM, higher plantar-flexor EMG amplitude at foot contact and early stance, and increased alpha and beta band intermuscular coherence at mid-stance compared to the control group. Second, imposing a mechanical knee constraint on healthy participants only increased beta-band intermuscular coherence between MG and LG muscles at the end of stance phase. Third, although the artificial KFC reproduced the sagittal knee posture observed in chronic KFC, the chronic KFC group still showed higher beta-band coherence across all plantar-flexor pairs, with additional increases in alpha-band coherence for the MG-LG and MG-SOL pairs and gamma-band coherence for the MG-LG pair. These results support our hypothesis that chronic KFC is associated with altered neural control of plantar flexors, and that the mechanical knee joint constraint reproduces only part of this altered pattern. These findings indicate that restricted knee motion alone cannot fully explain the neural organization of plantar-flexor activity in individuals with non-neurological knee flexion contracture.

### How is neural drive to the ankle plantar flexors altered in individuals with chronic KFC?

The chronic group exhibited altered neural drive to the plantar flexors, as evidenced by differences in both EMG amplitude and intermuscular coherence in the alpha and beta bands. In particular, the identification of an activation peak of plantar flexors occurring very early, near initial foot contact, indicates a modified motor control strategy during weight acceptance. Although this earlier peak pattern has been reported in neurological KFC, such as cerebral palsy (Romkes and Brunner, 2007), it has not been described previously in non-neurological chronic KFC. By contrast, studies in knee osteoarthritis and older adults have mainly reported increased or prolonged activation of lower-limb muscles, including the gastrocnemii during stance, rather than a clear shift of plantar-flexor peak activation toward initial contact (Hubley-Kozey et al., 2006; Rutherford et al., 2013; Schmitz et al., 2009). The increased plantar-flexor activation observed in the chronic group may reflect a compensatory neuromuscular strategy aimed at enhancing limb support, forward progression, and balance during stance (Lim et al., 2022), while also contributing to protective gait adaptations (Núñez-Cortés et al., 2023).

Regarding the increased intermuscular coherence observed in the chronic group, little is known about changes in common neural input to motor neurons during gait in chronic musculoskeletal impairments. Increased intermuscular coherence has been reported in patients with Parkinson’s disease (Flood et al., 2019), suggesting increased common synaptic input across multiple frequency bands, particularly in the beta band. In healthy individuals, increased intermuscular coherence has been attributed to joint stabilizing function during the cycling task (De Marchis et al., 2015). Consistent with this interpretation, Roeder et al. showed that intermuscular coherence during gait is strongly modulated within the gait cycle, with coherence peaking around heel strike and toe-off and largely absent during mid-stance, suggesting a role in phase-specific stabilization during locomotion (Roeder et al., 2024). This matched the temporal pattern we observed in the control group with and without an imposed knee constraint, whereas this temporal modulation was reduced in the chronic group revealing intermuscular coherence throughout the stance phase. During unipedal postural task, increased beta-band coherence has been attributed to compensation for smaller muscle mass in older adults (Nojima et al., 2020). Furthermore, beta and gamma bands intermuscular coherence appear to be increased during proprioceptive (perturbation-based) and proactive (obstacle-negotiation) locomotor tasks, suggesting greater cortical involvement (De Freitas et al., 2026). In contrast, during gait, beta–gamma intermuscular coherence in lower-limb synergists has been reported to be lower in older vs. young adults, which could be attributed to natural alterations in the corticospinal tract with age (De Freitas et al., 2026; Roeder et al., 2024). In stroke patients, a reduction in common corticospinal drive, accompanied by diminished coherence in the 10–25 Hz range has been found (Nielsen et al., 2008). Notably, older adults during walking have also been shown to exhibit increased beta-band coherence under fatigue, whereas young adults show no significant changes (Dos Santos et al., 2020). This increase with fatigue has been interpreted as a compensatory strategy to maintain neural control of ankle muscles and preserve motor output.

Taken together, the findings suggest that chronic KFC is associated with a fundamentally altered temporal pattern of plantar-flexor control during walking. In both control conditions, including walking with an artificial constraint, intermuscular coherence was concentrated around key gait events and diminished during mid-stance, despite the presence of substantial plantar-flexor activity. In contrast, individuals with chronic KFC demonstrated elevated EMG amplitudes at initial contact, accompanied by sustained intermuscular coherence during mid-stance. This divergence indicates that the chronic KFC pattern cannot be explained solely by altered joint mechanics or a flexed-knee gait posture. Rather, the concurrent persistence of muscle activation and common neural drive suggests a more continuously coupled control strategy (Boonstra et al., 2009b; Laine and Valero-Cuevas, 2017), whereby the plantar flexors remain coordinated across a larger portion of the stance phase instead of being synchronized primarily during gait transitions. Such a strategy may represent a long-term neuromuscular adaptation aimed at maintaining stability in the presence of chronic locomotor impairments. Future longitudinal studies are needed to determine whether this altered temporal organization of muscle activation and coordination can be modified through interventions targeting joint mechanics, gait retraining, or neuromuscular control.

### To what extent is this altered neural drive explained by the mechanical constraint at the knee?

The artificial knee constraint showed only a limited effect on plantar flexor neural control, particularly as an increase in beta-band intermuscular coherence between the biarticular plantar flexors (MG-LG pair). Artificial knee constraint has previously been used to simulate KFC in cerebral palsy and to examine its influence on lower-limb muscle synergies, including the total variance accounted for by one synergy as an index of neural control complexity (Cruz-Montecinos et al., 2021; Spomer et al., 2022). In healthy individuals, the simulated KFC did not alter muscle synergies, suggesting that mechanical restriction alone is insufficient to explain the neural control alterations reported in cerebral palsy (Spomer et al., 2022). The greater beta coherence observed in our study between the MG-LG pair may reflect a task-specific adaptation linked to the biarticular function of the gastrocnemii. This response may be related to increased shared neural drive and proprioceptive feedback during acute mechanical constraint, helping to reinforce plantar-flexor coordination and thereby contributing to ankle force regulation and limb stabilization during gait. In line with this interpretation, previous studies have reported that greater ankle postural demands may increase beta-band intermuscular coherence (Nojima et al., 2020; Watanabe et al., 2018; Yamanaka et al., 2023), supporting its role in the neural coordination of ankle musculature under mechanically challenging conditions (Jensen et al., 2018; Laine and Valero-Cuevas, 2017).

Notably, although the chronic and artificial groups showed similar knee kinematics, the chronic group exhibited greater plantar-flexor EMG amplitude during early stance and higher beta-band intermuscular coherence across all plantar-flexor pairs at mid-stance. This dissociation suggests that changes in neural control in the chronic condition cannot be explained solely by the mechanical constraint (Cruz-Montecinos et al., 2021). Several mechanisms may account for this difference. People with KFC have likely been exposed over time to the joint constraint, recurrent pain episodes, modified loading patterns, and compensatory gait strategies (Campbell et al., 2021; Cruz-Montecinos et al., 2022). Thus, the neural control pattern associated with long-term knee joint constraint may reflect an adaptive mode of neural control among synergistic muscles, rather than a purely biomechanical response to knee constraint. Although the artificial restriction model reproduced some aspects of the mechanical constraint, it lacked the chronic sensorimotor history present in the chronic group.

### Potential clinical implications

Traditional orthopedic approaches, such as total knee replacement, focus on correcting or compensating for the joint restriction and pain reduction to improve function. The present findings suggest that this mechanical perspective, while important, may not be enough to restore locomotion in people with non-neurological KFC. Interventions may also need to address the secondary developed neural impairment. This could include task-specific gait retraining (e.g., Functional Electrical Stimulation retraining or biofeedback) or explore tendon vibrations interventions aimed at restoring neuromuscular control (Marchand et al., 2025; Nojima et al., 2026; Parikh et al., 2025). Longitudinal studies are needed to explore these potential therapeutic alternatives.

### Strengths and Limitations

A key strength of this study is the implementation of an artificial constraint group, which effectively distinguishes acute mechanical restriction from chronic neuromuscular adaptation. However, several limitations should be also considered when interpreting this study. First, the chronic KFC group was small, which is understandable in a clinically specific population but nonetheless limits statistical power. Second, our study assessed the acute responses (1-2 minutes) to knee joint constraint and therefore cannot determine how the observed neuromuscular pattern develops over time. Third, although the artificial constraint was designed to simulate reduced knee motion, it cannot reproduce the full clinical reality of chronic haemophilic arthropathy, including pain history, structural joint changes, sensorimotor adaptation, and long-term behavioral compensation. Fourth, EMG amplitude was normalized to the peak amplitude of the signal rather than to MVC. This approach was chosen because MVC-based normalization may not be reliable in chronic group, given the potential of provoking pain and, hence, not obtaining true maximal force.

## Conclusions

Increased plantar-flexor activation and intermuscular coherence in the chronic constraint group compared to the artificial constraint group indicate that the limited ROM alone does not explain altered neural control of plantar flexors, suggesting central nervous system adaptations in individuals with non-neurological knee flexion contracture. These findings underscore the importance of targeting both joint mechanics and neural control in rehabilitation. Future longitudinal studies and approaches to assess cortico-muscular drive are needed to confirm these results.

## Supporting information

COMPLEMENTARY MATERIAL

## Credit authorship contribution statement

Carlos Cruz-Montecinos: Conceptualization, Methodology, Investigation, Formal analysis, Writing – original draft.

Tjeerd W. Boonstra: Methodology, Software, Writing – review & editing.

Huub Maas: Conceptualization, Supervision, Writing – review & editing.

## Declaration of Competing Interest

The authors declare no competing interests.

## Data Availability Statement

The datasets generated and analyzed during the current study are available from the corresponding author upon reasonable request.

## Funding

No specific funding was received for this study.

## Notes

### Competing Interest Statement

The authors have declared no competing interest.

