## Supplementary material for "The Limited Range of Motion of the Knee Does Not Fully Explain the Altered Neural Control of Plantar Flexors During Gait in Non-Neurological Knee Flexion Contracture": COMPLEMENTARY MATERIAL

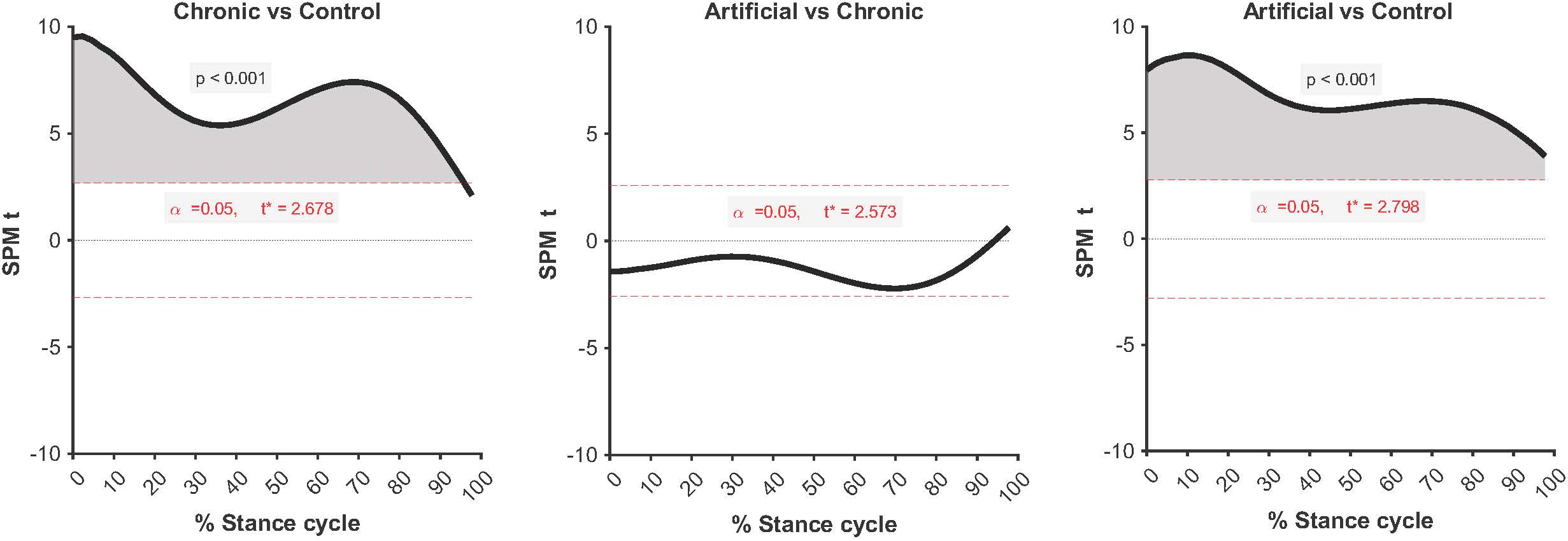
**Complementary Figure 1.** Statistical Parametric Mapping (SPM{t}) analysis of knee flexion-extension kinematics during the stance phase of gait. Panels show comparisons between chronic and control groups (left), chronic and artificial groups (middle), and paired comparisons between artificial and control conditions (right). Grey shaded regions indicate significant supra-threshold clusters (p < 0.05).

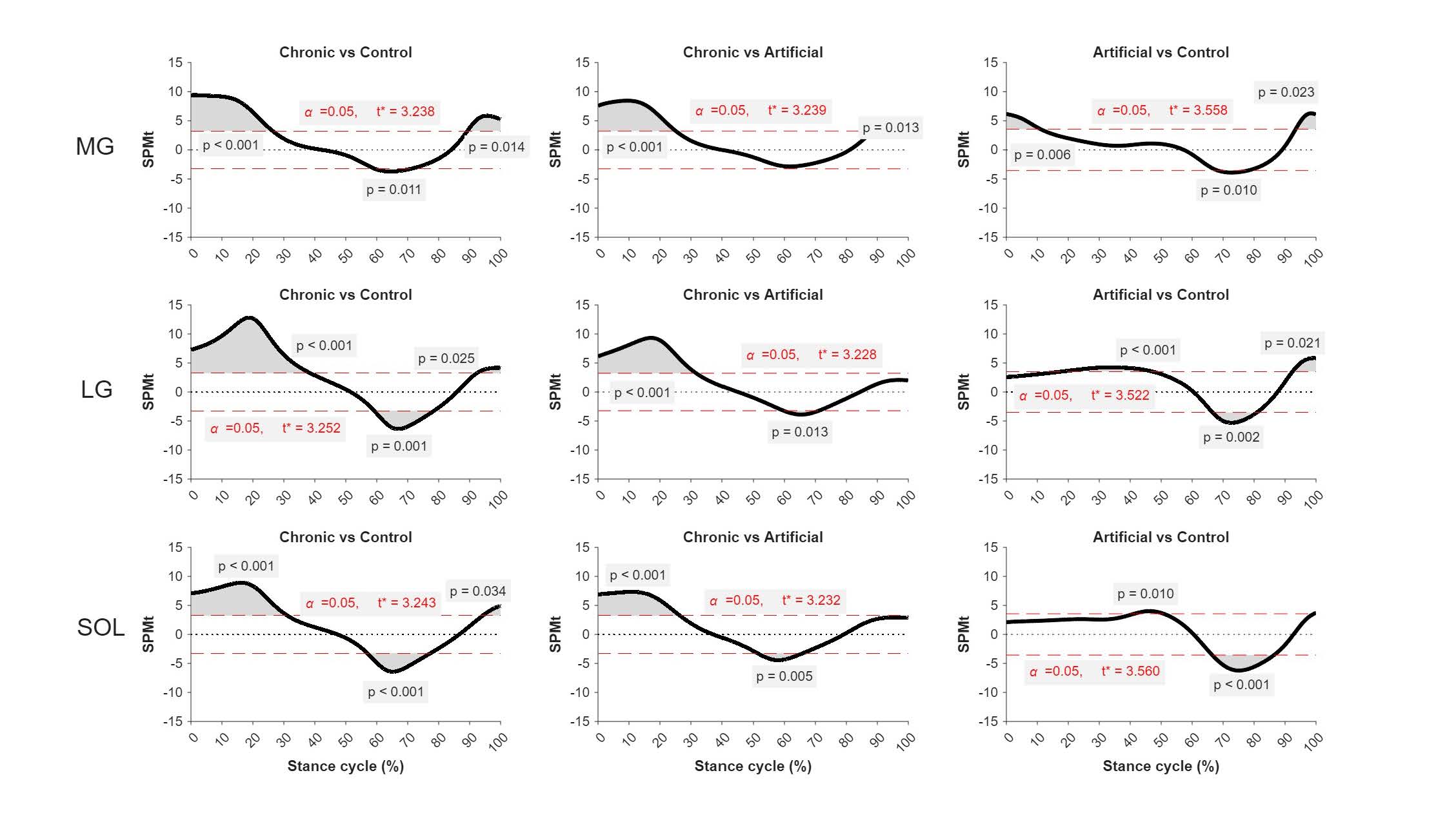

**Complementary Figure 2.** Statistical Parametric Mapping (SPM{t}) analysis of plantar-flexor EMG amplitude envelopes during the stance phase for the medial gastrocnemius (MG), lateral gastrocnemius (LG), and soleus (SOL) muscles. Panels show comparisons between chronic and control groups (left), chronic and artificial groups (middle), and paired comparisons between artificial and control conditions (right). Grey shaded regions indicate significant supra-threshold clusters (p < 0.05).
